# The impact of DNA extraction on the quantification of *Legionella*, with implications for ecological studies

**DOI:** 10.1101/2024.03.19.585788

**Authors:** Alessio Cavallaro, Marco Gabrielli, Frederik Hammes, William J. Rhoads

## Abstract

Monitoring the levels of opportunistic pathogens in drinking water is important to plan interventions and understand the ecological niches that allow them to proliferate. Quantitative PCR is an established alternative to culture methods that can provide a faster, higher throughput, and more precise enumeration of the bacteria in water samples. However, PCR-based methods are still not routinely applied for *Legionella* monitoring, and techniques such as DNA extraction differ notably between laboratories. Here, we quantify the impact that DNA extraction methods had on downstream PCR quantification and community sequencing. Through a community science campaign, we collected 50 water samples and corresponding shower hoses, and compared two commonly used DNA extraction methodologies to the same biofilm and water phase samples. The two methods showed clearly different extraction efficacies, which was reflected in both the quantity of DNA extracted and the concentrations of *Legionella* enumerated in both the matrices. Notably, one method resulted in higher enumeration in nearly all samples by about one order of magnitude and detected *Legionella* in 21 samples that remained undetected by the other method. 16S rRNA amplicon sequencing revealed that the relative abundance of individual taxa, including sequence variants of *Legionella*, significantly varied depending on the extraction method employed. Given the implications of these findings, we advocate for improvement in documentation of the performance of DNA extraction methods used in drinking water to detect and quantify *Legionella*, and characterise the associated microbial community.

## 1. Introduction

Accurate pathogen detection and quantification in drinking water is necessary to establish effective policies and minimise the infection risk. *Legionella* is a large genus of waterborne bacteria comprising more than 60 species, of which about half have been identified as opportunistic human pathogens (Fields *et al*., 2002). Current monitoring strategies heavily rely on culture-based methods (International Organization for Standardization, 2017), which have well documented limitations, including being time-consuming, producing less precise results, and having a bias towards *Legionella pneumophila* (Toplitsch *et al*., 2021). Molecular-based methods (i.e., quantitative/droplet digital qPCR/ddPCR) offer an alternative to culture that addresses some limitations associated with culture-based methods but have inherent limitations of their own. For example, molecular methods can be used to obtain more precise results faster, but detect genetic material from both live and dead organisms. While some legislation already allows for molecular methods to be used for compliance with monitoring requirements (European Parliament & European Council, 2020), the routine use of such methods remains a future prospect.

DNA extraction represents an essential step for molecular quantification of organisms that affects every downstream molecular analyses. DNA extraction methods consist of 1) lysis of cell membranes and nucleus in order to release DNA; 2) separation of DNA from other cellular components and debris; and 3) purification of the DNA. How well these steps are executed affects DNA yield, quantification of target organisms, and characterization of microbial community composition from various matrices (Martin-Laurent *et al*., 2001, Hart *et al*., 2015, Douglas *et al*., 2020, Cerca *et al*., 2022). Although all the DNA extraction methods share these general procedures, numerous commercial kits, protocols, and *ad-hoc* adaptations are available and described in literature. For example, protocols.io, a large repository of laboratory protocols contributed by researchers, has more than 2000 entries for DNA extraction methods and optimizations (https://www.protocols.io/search?q=%22dna%20extraction accessed on 01-12-2023). While different commercial kits and adaptations are usually implemented to extract DNA from specific matrices, thus not performing optimally on others, it is notable that there are examples where even different versions of the same commercial kit can produce different amount and quality of extracted DNA (Pearman *et al*., 2020). Despite the documented differences introduced by the DNA extraction can have on downstream analysis and results, no information on how different DNA extraction procedures influence the quantification of *Legionella* spp. is available to our knowledge.

There is generally a lack of adequate reporting of DNA extraction recoveries and potential biases in the drinking water field, as highlighted by the Environmental Microbiology Minimum Information (EMMI) guidelines (Borchardt *et al*., 2021). To address this, the EMMI guidelines propose the use of negative and positive controls to identify potential contaminants or issues with the DNA extraction process. An external positive control is particularly important to quantify extraction efficacy. The guidelines appropriately leave the specific choice of control to the study authors (Borchardt *et al*., 2021), but as a result, there is no established practice for *Legionella* DNA extraction from environmental samples. Using a pure culture, synthetic mixture of pure cultures, or cultures phagocytized by a host organism are all logical and common choices. However, these would provide little information about how well the extraction method extracts *Legionella* DNA from an environmental sample with a complex mixed microbial community, or about potential biases it introduces when performing sequencing analysis. From an ecological point of view, little information about the influence of DNA extraction on the microbial community characterisation in drinking water systems is available, particularly when wanting to distinguish between water and biofilm phases.

In this study, we process water and biofilm samples collected via a community science sampling campaign using two DNA extraction methods common in drinking water studies to demonstrate how the variability between and among methods can impact environmental sampling interpretation, with a specific focus on the quantification of *Legionella* spp. and community structure.

## 2. Materials and Methods

### 2.1 Field-scale community science sample collection and processing

#### 2.1.1. Participant recruitment and sampling

Employees from two research institutes were recruited via listserv email to collect shower water and biofilm samples from their homes. Participants received a pre-labelled sampling kit that contained a sterile 1 L glass bottle, a new shower hose to replace their existing one, sealing caps to retain the water within their harvested shower hose, and detailed sampling instructions. Briefly, following >8 h stagnation, the participant opened the shower tap and filled the 1 L bottle with the first flush of hot water until the bottle was full but not overflowing. The participant then removed the shower head and detached the existing shower hose, keeping the ends at approximately the same height to prevent water in the hose from leaking out. The used shower hose was capped with new threaded PVC caps to retain the water inside the hose and replaced with a new one. Samples were delivered to the lab for processing within 24 h. The experimental workflow is shown in Fig. 1.

**Figure 1.**
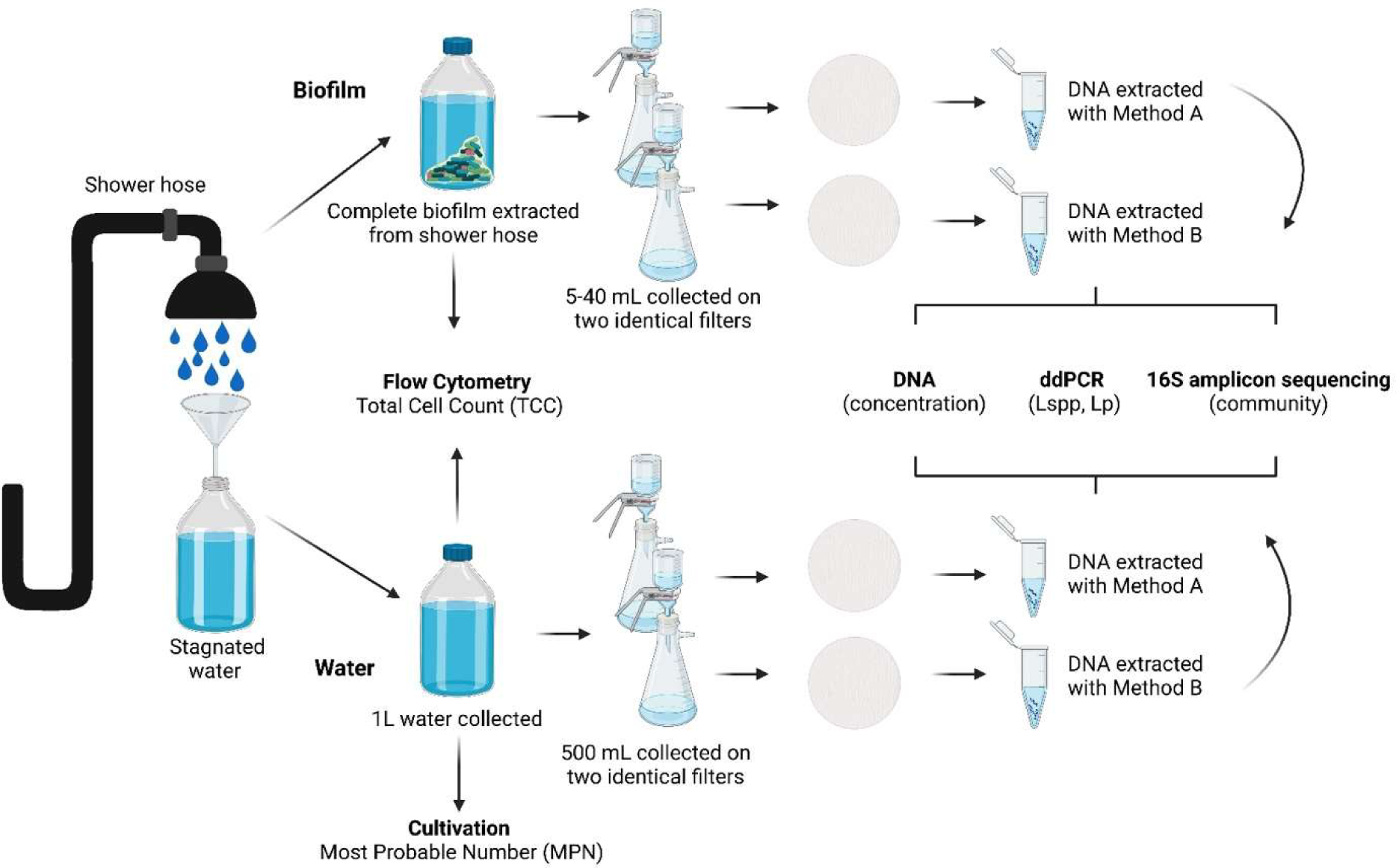
Overview of the experimental workflow used in this study. The shower hoses were collected with the first litre of 8-hours stagnated water. The biofilm was extracted from the shower hoses and then the biofilm and the water phase were: 1) used for flow cytometry to measure the total cell count (TCC): 2) filtered twice onto two different filters from which the DNA was extracted using two different commercially available kits.

#### 2.1.2. Sample processing - water

After mixing the 1 L sample, 100 mL was aliquoted to a sterile glass container, total cells quantified using flow cytometry, and *L. pneumophila* quantified using IDEXX’s Legiolert liquid culture kit according manufacturer instructions. The remains of the water sample were filter-concentrated onto duplicate 0.2 µm polycarbonate filters, recording the volume filtered, each filter was fragmented using a flame-sterilised scalpel and tweezers, and frozen at −20 °C until DNA extraction was carried out.

#### 2.1.3. Sample processing - biofilms

Water contained in the shower hose was collected and the hose exterior sheath (if applicable) was removed. The hose was filled with 20 mL of autoclaved 2 mm glass beads and biofilm was eluted into filter-sterilised and autoclaved water through five rounds filling the hose with the sterile water, sonication of the hose using the vibration of the beads to suspend attached biomass, and then collecting the water within the hose after each sonication. Each time the water was collected, the end from which the water flowed from the hose was alternated to create as much flow reversal and mixing as possible. After the last round of sonication/collection, the glass beads were removed, the hose filled with 50 mL water a final time, inverted 30 times, and water collected into the sample bottle. The total amount of water used for biofilm elution, and the length and diameter of the hose were recorded. Total cells were measured in the suspended biofilm sample after three rounds of 30 s sonication of a 2 mL aliquot at 40% amplitude and diluting 1:1,000 or 1:10,000 as necessary. Eluted biofilm from the sample bottle was filter-concentrated onto duplicate 0.2 µm polycarbonate filters until clogging (5-70 mL), each filter fragmented using a flame-sterilised scalpel, and frozen at −20 °C until DNA extraction was carried out.

#### 2.1.4. DNA extraction

Two commercially available DNA extraction kits with the same adaptation were used on each water and biofilm sample set. Each kit is commonly used for extracting DNA from environmental water samples and contains chemical (lysis solution) and physical (bead beating) cell lysis steps prior to column concentration and washing via centrifugation. Adaptations to the manufacturer instructions to each kit included submerging fragmented filters in 6 µL Lysozyme (50 mg/µL) and 294 µL 1X TE buffer for 1 hour at 37 °C mixing at 300 rpm; adding 30 µL Proteinase K (20 mg/mL) and 300 µL DNA extraction kit cell lysis solution, and continuing incubation for 30 minutes at 56 °C mixing at 300 RPM; and addition of 600 µL chloroform (isoamyl alcohol, 24:1 suitable for nucleic acid purification) with DNA extraction kit lysis beads. These adaptations were previously described by Voslo et al. (Vosloo *et al*., 2019). The beads, dissolved filters, and final 1230 µL solution were then bead-beaten on a vortex shaker at maximum speed for 5-minutes. Afterward, manufacturer instructions were followed. Each time DNA extraction was performed, a DNA extraction negative (un-used filter) and positive control was processed. The positive control consisted of replicate 100 mL bulk water samples collected from a bioreactor with colonized by native *Legionella* (Additional details provided in Supplementary Fig. 5). The bioreactor water was collected in bulk, then 100 mL aliquots concentrated onto replicate filters and frozen until DNA was extracted. *L. pneumophila* liquid culture and total cells were quantified in the bioreactor water to provide independent comparison of the extraction efficiency of *L. pneumophila* and total cells. The impact of each pre-treatment step on the DNA extraction was been evaluated and reported in Supplementary Fig. 1. Each kit contained steps to bind DNA followed by washing in an ethanol-based solution. Final elution volume was 100 µL.

### 2.2 Water quality measurements and methodology

#### 2.2.1. Total cells counts and extracted DNA

Total cells were quantified using a CytoFLEX (Beckman Coulter, Brea, USA) flow cytometer in 250 µL aliquots stained using SYBR® Green I (SG, Invitrogen AG, Basel, Switzerland; 10,000x diluted in Tris buffer, pH 8). Stained cells were incubated for 15 minutes at 37 °C prior to analysis (Prest *et al*., 2013). Extracted DNA was measured using Invitrogen Qubit dsDNA HS assay with a linear detection range of 0.2-100 ng double stranded DNA.

#### 2.2.2. Legionella spp. and L. pneumophila gene copy enumeration

*Legionella* spp. (ssrA) and *L. pneumophila* (mip) were measured with digital droplet polymerase chain reaction (ddPCR) using gene targets based on previously published assays validated to ISO SO TS12869:201 (Benitez & Winchell, 2013, Collins *et al*., 2017) and adapted for the ddPCR platform (Rhoads *et al*., 2022). Primer and probe sequences, master mix composition, and thermocycling conditions can be found in the supplemental material (Supplementary Table 1). A ddPCR reaction negative control (DNAse free water) was included for each batch of master mix prepared and was always negative. A ddPCR reaction positive control (Centre National de Référence des Légionelles) was included on each thermocycling run. Droplet formation and PCR thermocycling were performed using a Stilla geode and read using a Prism6 analyser with Crystal Reader software imaging settings pre-set and optimised for PerfeCTa multiplex master mix. Droplets were analysed using Crystal Miner software. Only wells with enough total and analysable droplets, as well as a limited number of saturated signals, were accepted according to Crystal Miner software quality control. Positive droplets were delineated using polygons, with positive wells being considered as those resulting in at least three droplets within the polygon. The limit of detection (LOD) was 5 gc/reaction (1 gc/µL template), and the limit of quantification (LOQ) was 12 gc/reaction (2.4 gc/µL template; Rhoads et al., 2022). Any sample with significant intermediate fluorescence clusters (i.e., “rain”) was diluted 1:10 and rerun.

### 2.3. 16S rRNA amplicon sequencing

For sequencing, the V4 region of the 16S rRNA gene was amplified by PCR using the primers Bakt_515F - Bakt_805R (Caporaso *et al*., 2011) and the DNA was quantified by Qubit dsDNA HS Assay. Samples were diluted, where possible, to the concentration of 1 ng/µL. A two – step PCR protocol was used to prepare the sequencing library: a first amplification (target PCR) was carried out with 1X KAPA HiFi HotStart DNA polymerase (Roche), 0.3 µM of each 16S primer and 2 µL of template DNA. After amplification, the PCR products were purified with the Agencourt AMPure System (Beckman Coulter, Inc.). The second PCR (adaptor PCR) was performed with limited cycles to attach specific sequencing Nextera v2 Index adapter (Illumina). After purification, the products were quantified and checked for correct length (bp) with the High Sensitivity D1000 ScreenTape system (Agilent 2200 TapeStation). Sample concentration was adjusted, and samples were subsequently pooled together in a library at a concentration 4nM. The Illumina MiSeq platform was used for pair-end 600 cycles (16S) with 10% PhiX (internal standard) in the sequencing run. Negative controls (PCR grade water) and a positive control (self-made MOCK community) were incorporated. Primer sequences, master mix composition, and reaction conditions can be found in the Supplementary Information (Supplementary Table 2). These steps were performed in collaboration with the Genetic Diversity Centre (GDC) of ETH Zurich.

### 2.4. Data analysis

16S rRNA sequencing data were processed on HPC Euler (ETHZ) using workflows established by the GDC (ETHZ, Zurich). Detailed data processing workflows are provided in the supplementary materials. For the 16S dataset, all R1 reads were trimmed (based on the error estimates) by 25 nt, the primer region removed, and quality filtered. Ultimately, sequences were denoised with error correction and chimera removal and amplicon sequence variants established using UNOISE3 (Edgar & Flyvbjerg, 2015). In this study, the predicted biological sequences will be referred to as Zero-Radius Operational Taxonomic Units (ZOTUs). Taxonomic assignment was performed using the Silva 16S database (v128) in combination with the SINTAX classifier. Samples were not rarefied to avoid the loss of data due to differences in the sequencing depth (McMurdie & Holmes, 2014). Distance ordination and relative abundance were calculated using R (version 4.2.1) and R studio (version 2022.07.2+576) using the Bioconductor package “phyloseq” (version 1.42.0, (McMurdie & Holmes, 2013). LefSe analysis was performed using Microbiome Analyst (Chong *et al*., 2020). SparCC correlation analysis was performed using the software FastSpar (Watts *et al*., 2019). All graphs were constructed with the R package “ggplot2” (version 3.4.0). Unless otherwise specified, all packages were operated using the default settings. Absolute abundance for *Legionella* quantification was calculated as follows:

Absolute Abundance = Relative Abundance (16S amplicon sequencing) X Total Cell Count (Flow Cytometry) X DNA extraction efficacy

### 2.5. Data availability

The data for this study have been deposited in the European Nucleotide Archive (ENA) at EMBL-EBI under accession number PRJEB72629. All other datasets from this study is available from the Eawag Research Data Institutional Repository.

## 3. Results

### 3.1. Comparison of DNA extraction efficacies

The two DNA extraction methods used in this study yielded different amounts of DNA (Fig. 2). Method A extracted substantially more DNA overall (median value 1.03 fg/cell, IQR: 1.70; n = 83) than method B (median value 0.03 fg/cell, IQR: 0.06; n = 74) (Wilcox paired test - p-value: < 0.001). The median DNA/cell value detected with method A was 186-fold higher than that of method B for water samples, but only 13-fold higher for biofilm samples. However, this could partially be artificial due to the fact that biofilm samples seem to reach a plateau at about 10^4^ ng of extracted DNA (see section 4.2). Method B failed to extract detectable levels of DNA from 15 out of 50 water samples. These samples with low/no-extracted DNA were still used for *Legionella* quantification through ddPCR, but were excluded from the sequencing run.

**Figure 2.**
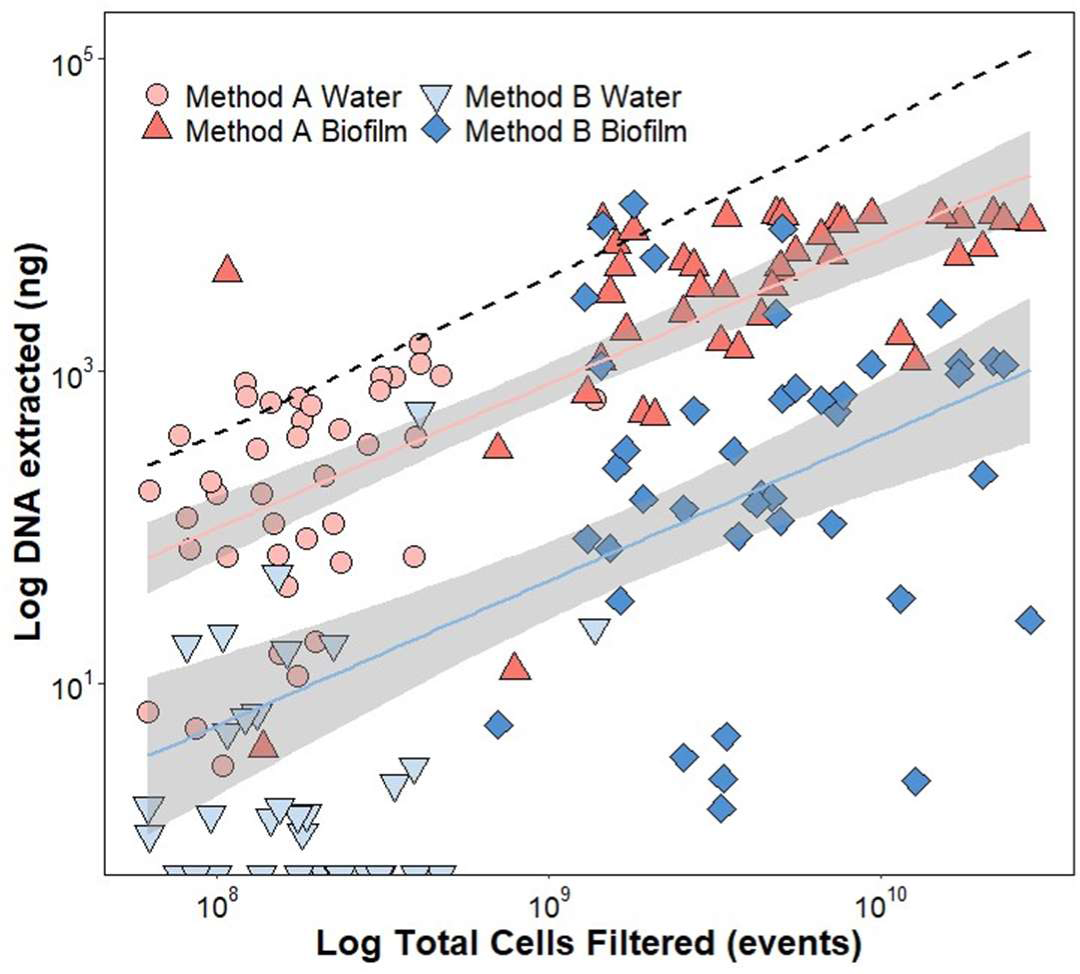
DNA extraction efficacy of two tested extraction methods on the same water and biofilm samples collected from shower hoses in private homes. Method A samples are red, method B samples are blue. The samples shown in lighter colour refer to the water phase, while the darker colour indicates the biofilm phase. The dashed line represents a theoretical maximum DNA extraction efficacy of 4 fg/cell. Markers crossing the axes represent samples with no quantifiable extracted DNA.

To calculate the extraction efficacy for each sample, an average DNA-per-cell value of 4 fg/cell was used (Christensen *et al*., 1995). The median value of the extraction efficacy for the water samples extracted with method A was 35 %, while it was 0.2 % for method B. For biofilm samples, the median value was 21.7 % for the ones extracted with method A and 1.6 % for method B. A few samples had an extraction efficacy above 100%, which is likely because the chosen value of 4 fg/cell does not consider the heterogeneity in DNA content across different bacteria, particularly in complex communities (Button & Robertson, 2001).

### 3.2. Impact on *Legionella* quantification

The differences in the DNA extraction affected the detection and quantification of *Legionella* spp. and *L. pneumophila*. The quantification of *Legionella* spp. using ddPCR was significantly higher (Wilcox paired test – p-value < 0.001) for the samples extracted with method A than the ones extracted with method B, showing a 39-fold median increase of gene copies detected per 100 mL (Fig. 3a). *Legionella* spp. was not detected with ddPCR in 21 water samples from method B and 10 samples (one water and nine biofilm) from method A (Fig. 3a), but in all cases where *Legionella* spp. was not detected, it was detected in the same samples with the other method. In nine samples among the 21 non-detected by method B, there was not quantifiable DNA in the first place (section 3.1). As with *Legionella* spp., the quantification of *L. pneumophila* using ddPCR was significantly higher Wilcox paired test – p-value < 0.001) for the samples extracted with method A than the ones extracted with method B, with a 44-fold increase of gene copies per 100 mL detected (Supplementary Fig. 2).

**Figure 3.**
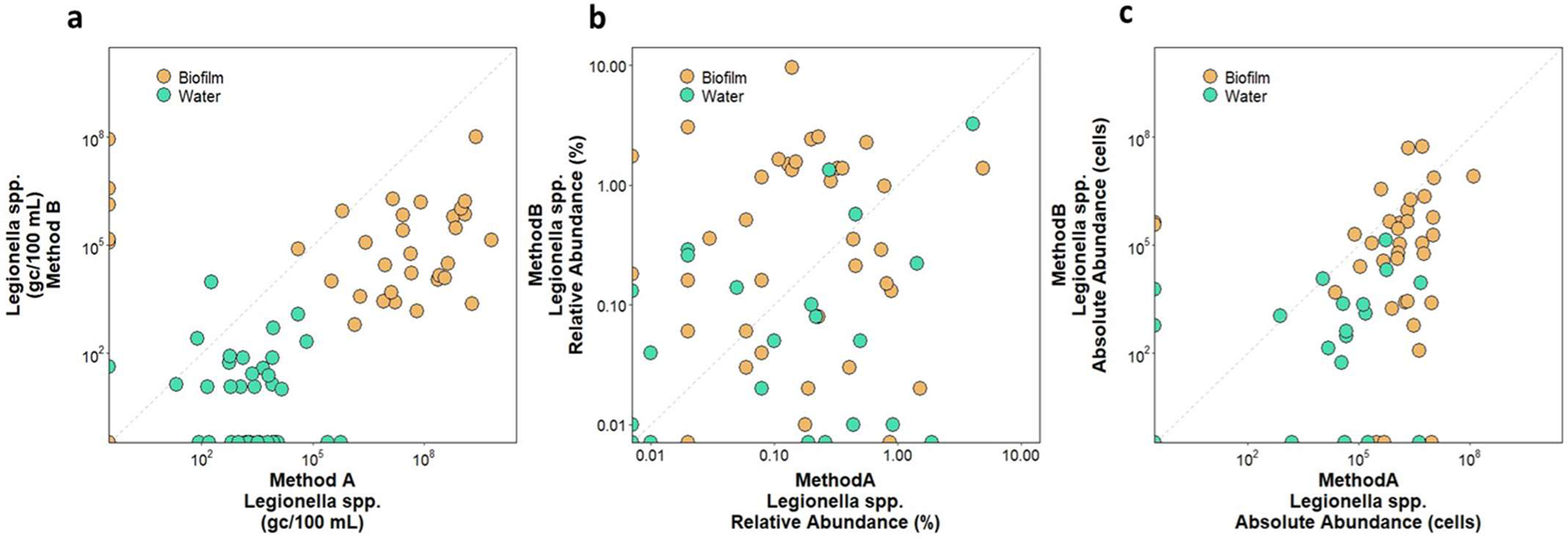
Comparison of two DNA extraction methods for the quantification of *Legionella* spp. (a) Number of gene copies per 100 mL using ddPCR; **(b)** Relative abundance of *Legionella* spp. calculated with 16S amplicon sequencing data; **(c)** Absolute abundance of *Legionella* spp. calculated by multiplying the relative abundance (shown in b) with the total number of filtered cells and the DNA extraction efficacy (Fig. 2). Biofilm samples are indicated in orange, while water samples are shown in green. Markers crossing the axes represent samples with no detected *Legionella* spp.

Overall, there was a 50% and 45% of presence-absence agreement between ddPCR and cultivation in the samples for method A and B, respectively (Supplementary Fig. 3). As expected, there instances when both methods detected *L. pneumophila* DNA but the cultivation data was negative. Method A had 16 samples that were positive by ddPCR but negative by culture (n=43, 37%); method B had 10 (n=42, 24%). However, there were also samples positive by culture but not by ddPCR. Method A had six samples that were culture positive but ddPCR negative (13%) while method B had 13 (31%). While this could be attributed to low DNA extraction efficacy for the 13 samples extracted with method B, the 6 samples extracted with method A showed overall good recovery (DNA extraction efficacy >50%), which cannot justify this observation.

When determining the relative abundance of *Legionella* spp. (Fig. 3b) from sequencing data, highly variable results were observed between methods. *Legionella* spp. was not detected through amplicon sequencing in six samples (four water, two biofilm) extracted with method B, and in four samples (two water, two biofilm) extracted with method A, and one sample with both methods. For a direct quantitative comparison with the ddPCR data, absolute abundance was calculated by multiplying by the TCC, while accounting for the DNA extraction efficacy (Fig. 3c). Quantification of *Legionella* spp. using amplicon-sequencing-derived absolute abundance remained statistically higher for method A than method B, showing a 50-fold difference (Wilcox paired test – p-value: < 0.001).

### 3.3. Impact on ecological observations

Besides *Legionella* quantification, the differences in the extraction methods also affected the community structure detected in the samples: bacterial genera were detected with different abundances in samples extracted with either method A or method B. The distance between samples was calculated using the Bray-Curtis index, which resulted in an NMDS plot with a stress value of 0.222 (Fig. 4). The analysis showed more significant clustering according to the phase of the samples (i.e., water or biofilm; PERMANOVA, p-value < 0.001, R2= 0.03), though the samples also cluster significantly with respect to the two DNA extraction methods used (PERMANOVA, p-value < 0.001, R2= 0.02). This suggests the selection of extract kit changes the community composition.

**Figure 4.**
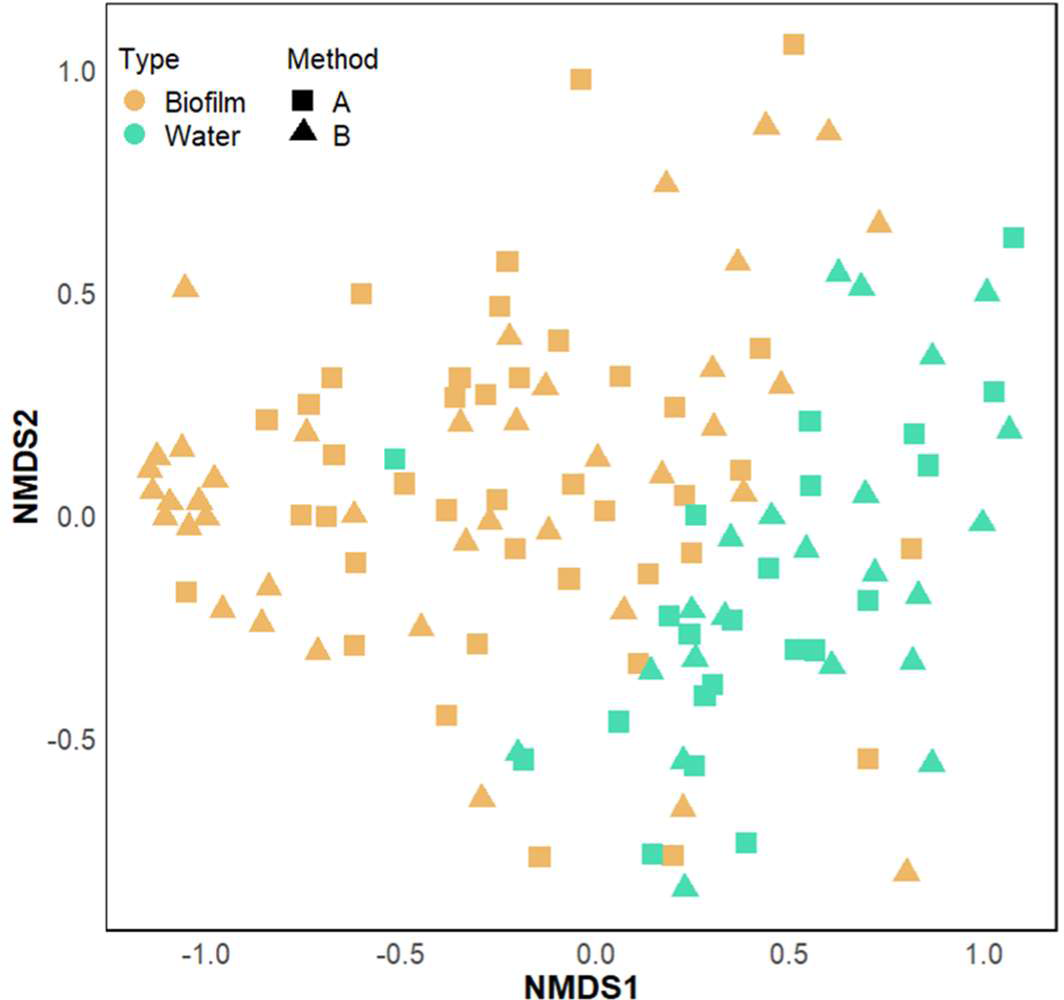
Ordination analysis of the 16S amplicon sequencing data, stress = 0.222. The graph shows distance among samples calculated using the Bray-Curtis method and plotted on a NMDS plot. The colour of the points refers to the phase (biofilm in orange; water in green), while the shape shows the method used to extract the DNA of the samples (Method A indicated with squares, method B indicated with triangles).

In order to provide a better overview of the ecological differences caused by the different DNA extraction methods, LefSe analysis was used to determine genera most likely to explain differences between the two (Fig. 5) (Segata et al. 2011). The analysis shows that the relative abundance of 52 genera are enriched in either method A (32 genera) or method B (20 genera) with LDA scores ranging between 2 and 5. Based on the effect size, we detected the genera *Sphingomonas*, *Thermus* and *Gemmata* to be the most enriched in method A (LDA scores: 5, 4.81, 4.59), while *Caulobacter*, *Obscuribacteraceae* and *Hirschia* were found to be the genera with a stronger effect size for method B (LDA scores: 5.05, 4.87, 4.52) (Fig. 5a).

**Figure 5.**
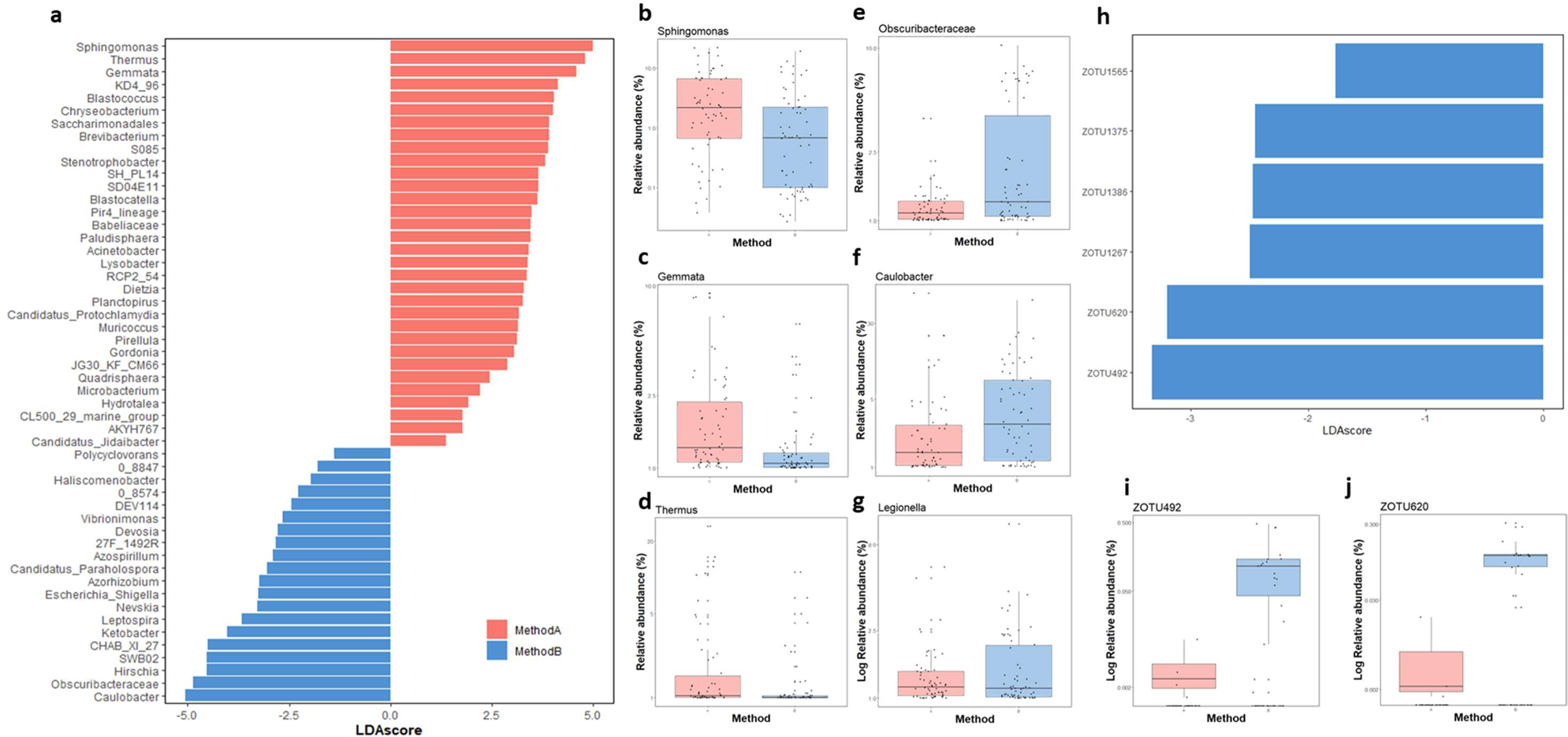
Differential abundance of bacterial genera and *Legionella*-associated zOTUs between the methods A (in red) and method B (in blue). A) LDA scores derived from the LefSe analysis showing the genera that are overall enriched in one method; B-D) Box plots of the relative abundance obtained using the two methods of the genera with the highest absolute LDA scores in Fig. 5a; G) Box plot for the relative abundance of *Legionella*; H) LefSe analysis showing the LDA scores of *Legionella*-associated-zOTUs that are enriched in either methods; I-J) Box plot of the relative abundances of the two *Legionella*-associated-ZOTUs with the highest absolute LDA scores in Fig. 5h.

While the genus *Legionella* was not detected by this statistical analysis as enriched in one specific method (Fig. 5g), a LefSe analysis performed at the zOTU level was able to detect 6 *Legionella*-associated-zOTUs (zOTU492, zOTU620, zOTU1267, zOTU1386, zOTU1375, zOTU1565) whose relative abundances were all enriched in method B (LDA score ranging 1.77 to 3.33), but not in method A. In one case, a *Legionella* zOTU (zOTU1267) was classified to the species level, and assigned to *L. geestiana*, while the other five zOTUs only had a classification to the genus level. SparCC analysis was performed to detect correlations between the zOTUs linked to the genus *Legionella* and the ones representing the rest of the community. For the biofilm samples extracted with method A, 53 significant correlations (correlation coefficient > 0.4) were detected (n samples = 45), while 3442 correlations were detected in the samples extracted with method B (n = 45). For the water phase, 83 significant correlations were detected in the samples extracted with method A (n = 39), while 225 were detected in the ones extracted with method B (n = 22). Among the correlations detected, five were shared between the methods in the biofilm phase, while 35 were shared in the water samples.

## 4. Discussion

### 4.1. The choice of the DNA extraction method is important

DNA extraction is a crucial step for downstream molecular analysis and has been identified as one of the most problematic steps, since it is prone to introduce biases and inefficiencies in extracting DNA (Costea *et al*., 2017, Brandt & Albertsen, 2018). Ineffective DNA extraction with lower DNA yields (Fig. 2) results in lower ddPCR concentrations for specific target organisms (Fig. 3a). This was previously demonstrated by Cerca and colleagues (Cerca *et al*., 2022), showing that DNA extraction efficacy is likely to influence the quantification of target organisms by qPCR in polymicrobial consortia. The authors extracted gDNA from three microorganisms in pure cultures at known concentrations (individually or collectively cultured) and performed qPCR generating different calibration curves, of which only the one that corrected for the DNA loss during the DNA extraction provided reliable results.

Several previous ecology studies noted the potential influence of DNA extraction methods on DNA yield, bacterial diversity and relative abundance measurements, for example in soil samples (Martin-Laurent *et al*., 2001), human breast milk (Douglas *et al*., 2020) and faecal samples (Hart *et al*., 2015). However, only a few studies have so far investigated the impacts of different DNA extraction methods on drinking water samples (Hwang *et al*., 2012, Brandt & Albertsen, 2018). While each of these studies identified the method that fit the need of the proposed experimental design, a perfect universal DNA extraction method does not exist at present, and the efficacy of the extraction protocol applied is dependent on the sample type or presence of specific organisms (Costea *et al*., 2017, Hermans *et al*., 2018, Brauer & Bengtsson, 2022). While these methodological differences are not surprising *per se*, little data is available on how DNA extraction influences detection and quantification of microorganisms in drinking water, how this specifically affects the quantification of the opportunistic pathogenic *Legionella* species, and how these challenges/problems can be mitigated if molecular methods are to be used quantitatively for routine monitoring within legislative frameworks.

We found that different methods extracted significantly different amounts of DNA from the same samples (Fig. 2), and that this consequently influenced the quantification of *Legionella* via ddPCR (Fig. 3a). We furthermore showed the extraction efficacy relative to an ideal situation in which all cellular DNA is extracted, using an average estimation of cellular DNA content (Fig. 2; (Christensen *et al*., 1995). However, this is an approximation that does not account for the variations in DNA content among different bacteria and requires access to a flow cytometry to obtain the total cell count. Other studies report the addition of external spike-in sequences to measure recovery of a known amount of target (Stoeckel *et al*., 2009). This approach does not provide, however, any information about the correct lysis of the cells. The latter can be achieved by spiking whole-cell-microorganisms at a defined amount, which provides quantifiable recovery of a target organism (or surrogate microbes) and is an approach also advocated for in the EMMI guidelines (Borchardt *et al*., 2021). Regardless of the approach used, here we advocate for consistent measurement and reporting of the extraction efficacy and the way it has been calculated, particularly in studies where quantitative data is generated. Such information allows an estimation of the DNA loss, and the potential for a corrective factor to be applied to quantitative data.

### 4.2. Water and biofilm phase

The water and biofilm phases revealed interesting differences in terms of DNA extraction. Both methods reported a lower DNA yield in DNA-per-cell for the water samples compared to the paired biofilm samples, but this effect was more pronounced for method B, which failed to extract detectable DNA from multiple water samples (Fig. 2). While the lower yield can be partially attributed to the presence of extracellular DNA in the biofilm matrix (Di Martino, 2018), the poor extraction efficacy recorded overall with method B can only be attributed to the weak performance of the method itself. We observed a maximum DNA recovery of approximately 10^4^ ng, which appears to be an artificial limit at high cell concentrations (Figure 2). One possible explanation for this could be that there is a maximum amount of nucleic acid that can bind to the extraction column used, which is referred to as binding capacity. While every kit has a distinct binding capacity, the plateau observed in our results is consistent to that reported by several manufacturers, and this can potentially lead to an underestimation of DNA concentrations and, by consequence, of specific target organisms when processing high biomass samples.

Our study also revealed a clear separation in the microbial communities detected in the water and in the biofilm phase, irrespective of the extraction method that was used (Fig. 4). This is an important ecological observation and only a few other papers have previously reported that different communities inhabit the two phases in a distinctive way. In an ecological investigation of the communities in an unchlorinated distributions system, Liu and colleagues (Liu *et al*., 2018) showed that bulk water and pipe biofilm clustered differently in a principal component ordination analysis, identifying key species dominating either the water (e.g. *Polaromonas* spp.) or the biofilm (e.g. *Pseudomonas* spp; *Sphingomonas* spp.). Similarly, Proctor and colleagues observed that the biofilm and bulk water bacterial communities were significantly different in shower hoses (Proctor *et al*., 2018). In particular, it was shown that three taxa accounting for 91% of the biofilm sequences only accounted for 31% of the cold-water sequences detected, while seven predominant taxa in the cold water accounted only for 2% of the biofilm sequences detected. One possible explanation for this phenomenon can be attributed to early establishment of defined ecological niches within the biofilm compared to higher dynamics of bulk water due to flow and exchange of nutrients. Biofilms have lower richness than the bulk water, meaning that only a few species dominate the biofilm ecological niches (Henne *et al*., 2012). This suggests that there is relatively low ability for transient bulk water organisms to establish in developed biofilms. This could also explain why it has been observed that the communities of biofilms are more stable in time that the ones present in water (Inkinen *et al*., 2014).

With respect to *Legionella*, these differences are important as it is known that the pathogen live at higher concentrations in biofilms (Declerck, 2010, Cavallaro *et al*., 2023), as is the case with most opportunistic building plumbing pathogens (Falkinham *et al*., 2015). We observed this phase-difference despite the fact that all shower hoses in our experiments were collected after stagnation, thus perhaps providing more opportunities for an exchange between the two phases. However, the result could partially be attributed to the sampling strategy adopted: we collected 1L water samples, thus not only the water that was stagnating in the shower hose (ca 80 mL). The additional water coming from building plumbing apart from the shower hoses might have masked any exchange between the two phases to some extent, thus increasing the magnitude of the separation observed (Fig. 4). This highlights the importance of understanding and correctly reporting all upstream processing steps that may influence downstream analysis; it also demonstrates the importance of how considering both phases in ecological studies and routine monitoring can provide more information than typical monitoring plans. Previous studies have already adopted strategies for on-site removal and replacement of portions of biofilm-containing-pipes, through dedicated points of entrance and aseptic insertion of new coupons (Deines *et al*., 2010). However, it is not possible to establish how small portions can be representative of the entire distribution network biofilm, which is an issue that must be further investigated.

### 4.3. Relative and absolute abundance

To quantitatively compare the abundances of microorganisms across different samples and studies analysed with 16S amplicon sequencing, it is useful to convert the relative abundances values into absolute abundance (Props *et al*., 2017). Several studies have reported the potential occurrence of opportunistic pathogens in environmental samples using sequencing data in the form of relative abundance (Neu *et al*., 2019, Paranjape *et al*., 2020, Cavallaro *et al*., 2023). While these observations are ecologically relevant, the use of relative abundance does not provide quantitative information on concentrations. An estimation of the absolute abundance can be obtained using different methods, of which the most used are (1) qPCR quantification of the total 16S rRNA gene and (2) flow cytometric total cell counts, chosen for this study (Galazzo *et al*., 2020, Wang *et al*., 2021). Some considerations are, however, necessary. Since both qPCR and sequencing use the same extract that is subjected to the same methodological biases, calculating the absolute abundance with 16S qPCR data has the downside of carrying any DNA loss occurred during the extraction, leading to the generation of inaccurate results. In this case, it is therefore necessary to account for the DNA loss during the extraction (i.e., determining the DNA extraction efficacy) for the calculation of the absolute abundance. Moreover, calculating the absolute abundance with qPCR harbours additional limitations: previous studies have in fact observed that qPCR can detect only large changes in gene concentrations and is strongly influenced by the primer pairs and the reaction conditions (Props *et al*., 2017). The quantification using flow cytometry, on the contrary, can overcome these limitations, as it is not necessary to calculate the DNA extraction efficacy separately in order to obtain the absolute abundance.

### 4.4. Relevance of an accurate *Legionella* detection and quantification

The use of different DNA extraction methods clearly has variable outcomes in relation to the accurate detection and quantification of target organisms (in this case *Legionella* spp.) in drinking water systems. This is relevant from two perspectives: 1) the implications for microbial ecology studies; 2) the implications for the monitoring of *Legionella* spp. under regulatory settings.

#### 4.4.1. Ecology

Ecological studies often work towards providing insights into the microbial composition of given samples under specific conditions, but an accurate detection of the organisms involved, and their relative proportions are crucial to define valuable biological interpretations of the data collected. The most common reported examples of how DNA extraction affects ecological observations are related to the studies conducted on the human microbiome. For example, in a study aiming at establishing the impact of DNA extraction procedures on the assessment of human gut composition, Kennedy and colleagues demonstrated that, within individual patients, community structures clustered together based on the kit used to extract the DNA from the samples (Kennedy *et al*., 2014). This effect was even more pronounced depending on the distance metrics that were used, and this had an important value given that the study was aimed at showing differences in community composition between volunteers and patients with inflammatory bowel disease (Kennedy *et al*., 2014). In a different study involving the human oral microbiome, Lazarevic and colleagues found that while the most abundant taxa were detected in the samples extracted with both methods, for some genera the relative abundances were significantly different depending on the kit used (Lazarevic *et al*., 2013). Similar results were obtained in a study that compared DNA extraction kits and primer sets for freshwater sediments samples: no significant diversity in terms of community structure was in fact detected, but the relative abundance of specific taxa varied significantly (Shi *et al*., 2022). Moreover, this study also highlighted differences in richness and relative abundance for the eukaryotic communities detected in samples extracted with two different kits.

In the context of the microbial ecology of *Legionella,* the relevance of this is mainly linked to the observational studies aiming at understanding how this opportunistic pathogen lives in aquatic systems through their relationship with the surrounding organisms (Paranjape *et al*., 2020, Paranjape *et al*., 2020, Scaturro *et al*., 2022). In a previous paper from our group, for example, we observed that *Legionella* spp. correlated positively and negatively with prokaryotic and eukaryotic microorganisms in biofilm samples from plumbing systems (Cavallaro *et al*., 2023). These studies used different combinations of statistical approaches that process the sequencing outcome (often in the form of relative abundance) to infer correlations. In this context, biases introduced by the DNA extraction (in terms of community structure, abundance and amplicon sequence variants detected) can influence the analysis leading to wrong ecological interpretations. Our results, for example, demonstrate that the correlations inferred with SparCC when using the sequencing data of samples extracted with different methods produce variable outcomes, both in the number of correlations and identity of the zOTUs involved. We argue that a correct understanding of the microbial ecology is crucial for the control of the pathogen in drinking water systems, and the possibility of comparing studies is an important tool towards this goal; however, biases due to the molecular protocol applied (i.e., DNA extraction) can work against the formulation of correct ecological hypothesis.

#### 4.4.2. Legislative compliance and risk assessment

A reliable quantification of *Legionella* spp. is important to accurately control the level of the pathogen in engineered aquatic systems and to assess the risk linked to its presence. Quantitative microbial risk assessment (QMRA) uses information regarding pathogen concentration to determine health implications of microbial hazards (Hamilton & Haas, 2016). Thus, the concentrations of the pathogenic organism investigated are of extreme importance to establish the risk associated with its exposure. With respect to QMRA of *Legionella*, most studies use concentrations measured with conventional culture approaches, which likely underestimates actual concentrations and, in turn, may lead to the underestimation of risk (Hamilton & Haas, 2016). Molecular methods (including qPCR or ddPCR) are generally more sensitive and overcome some of the limitations of traditional culture approaches, but, as demonstrated in this study, are subject to errors arising from DNA extraction efficacy differences. Therefore, the future inclusion of molecular methods in QMRA requires careful consideration, documentation and reporting of the entire sampling processing pipeline.

Similarly, the effects of the DNA extraction on the quantification of *Legionella* can also have an impact on the routine monitoring for the presence of *Legionella*. Normally, the water phase is collected, filtered, and then plated onto BCYE agar plates for enumeration and culture confirmation (International Organization for Standardization, 2017). Interventions are required when the concentration of *Legionella* reaches the threshold indicated by the national authorities (in Switzerland, 1000 CFU/L, (Eidgenössische Departement des Innern (EDI), 2016). This entire process, however, is time- and labour-consuming as culturing *Legionella* typically takes 7 - 14 days. Moreover, culturing does not account for bacteria in viable-but-not-cultivable (VBNC) state (Ramamurthy *et al*., 2014). Therefore, the demand for the implementation of molecular methods (i.e., quantification of *Legionella* through qPCR/ddPCR) in the context of assessing water quality has increasingly spread across practitioners and authorities and in fact, for example, the new EU legislation allows for alternative methods to be used (European Parliament & European Council, 2020). The implementation of such molecular methods in routine monitoring calls for a more detailed and standardized reporting of the protocols used.

Our data not only demonstrates the variability in terms of concentration of *Legionella* when using different DNA extraction methods, but it also highlights how the pathogen is not detected in some samples extracted with one method, while it is quantified when the DNA is extracted with the other method used in this study. This would obviously influence the legal settings, since a wrong estimation of the concentration of *Legionella* due to extraction biases can lead to either unnecessary interventions (which are expensive for the practitioners) or increased risks. While being aware of the challenges involved in the standardization of methods, we strongly believe that an extensive reporting of the protocol details (e.g., DNA extraction method used, extraction efficacy and how it was calculated) would be beneficial for a reliable comparison among studies, monitoring strategies and regulations.

## Supporting information

16S amplicon sequencing data preparation

## Acknowledgements

This research was funded by the Federal Food Safety and Veterinary Office (FSVO), in partnership with the Federal Offices of Public Health (FOPH) and Energy (SFOE) in Switzerland, through the project LeCo (Legionella Control in Buildings; Aramis nr.:4.20.01) and Eawag discretionary funding. Data produced and analyzed in this paper were generated in collaboration with the Genetic Diversity Centre (GDC), ETH Zurich.

## References

Benitez AJ & Winchell JM (2013) Clinical application of a multiplex real-time PCR assay for simultaneous detection of Legionella species, Legionella pneumophila, and Legionella pneumophila serogroup 1. J Clin Microbiol 51: 348–351.

Borchardt MA, Boehm AB, Salit M, Spencer SK, Wigginton KR & Noble RT (2021) The Environmental Microbiology Minimum Information (EMMI) Guidelines: qPCR and dPCR Quality and Reporting for Environmental Microbiology. Environ Sci Technol 55: 10210–10223.

Brandt J & Albertsen M (2018) Investigation of Detection Limits and the Influence of DNA Extraction and Primer Choice on the Observed Microbial Communities in Drinking Water Samples Using 16S rRNA Gene Amplicon Sequencing. Front Microbiol 9: 2140.

Brauer A & Bengtsson MM (2022) DNA extraction bias is more pronounced for microbial eukaryotes than for prokaryotes. Microbiologyopen 11: e1323.

Caporaso JG, Lauber CL, Walters WA, Berg-Lyons D, Lozupone CA, Turnbaugh PJ, Fierer N & Knight R (2011) Global patterns of 16S rRNA diversity at a depth of millions of sequences per sample. Proc Natl Acad Sci U S A 108 Suppl 1: 4516–4522.

Cavallaro A, Rhoads WJ, Sylvestre E, Marti T, Walser JC & Hammes F (2023) Legionella relative abundance in shower hose biofilms is associated with specific microbiome members. FEMS Microbes 4: xtad016.

Cerca N, Lima A & Franca A (2022) Accurate qPCR quantification in polymicrobial communities requires assessment of gDNA extraction efficiency. J Microbiol Methods 194: 106421.

Chong J, Liu P, Zhou G & Xia J (2020) Using MicrobiomeAnalyst for comprehensive statistical, functional, and meta-analysis of microbiome data. Nat Protoc 15: 799–821.

Christensen H, Olsen RA & Bakken RL (1995) Flow cytometric measurements of cell volumes and DNA contents during culture of indigenous soil bacteria. Microbial Ecology 29: 49–62.

Collins S, Stevenson D, Bennett A & Walker J (2017) Occurrence of Legionella in UK household showers. Int J Hyg Environ Health 220: 401–406.

Costea PI, Zeller G, Sunagawa S, et al. (2017) Towards standards for human fecal sample processing in metagenomic studies. Nat Biotechnol 35: 1069–1076.

Declerck P (2010) Biofilms: the environmental playground of Legionella pneumophila. Environ Microbiol 12: 557–566.

Deines P, Sekar R, Husband PS, Boxall JB, Osborn AM & Biggs CA (2010) A new coupon design for simultaneous analysis of in situ microbial biofilm formation and community structure in drinking water distribution systems. Appl Microbiol Biotechnol 87: 749–756.

Di Martino P (2018) Extracellular polymeric substances, a key element in understanding biofilm phenotype. AIMS Microbiol 4: 274–288.

Douglas CA, Ivey KL, Papanicolas LE, Best KP, Muhlhausler BS & Rogers GB (2020) DNA extraction approaches substantially influence the assessment of the human breast milk microbiome. Sci Rep 10: 123.

Edgar RC & Flyvbjerg H (2015) Error filtering, pair assembly and error correction for next-generation sequencing reads. Bioinformatics 31: 3476–3482.

Eidgenössische Departement des Innern (EDI) (2016) Verordnung des EDI über Trinkwasser sowie Wasser in öffentlich zugänglichen Bädern und Duschanlagen. p.^pp.

European Parliament & European Council (2020) Directive (EU) 202/2184 of the European Parliament and of the Council on the quality of water intended for human consumption. p.^pp.

Falkinham JO, 3rd, Hilborn ED, Arduino MJ, Pruden A & Edwards MA (2015) Epidemiology and Ecology of Opportunistic Premise Plumbing Pathogens: Legionella pneumophila, Mycobacterium avium, and Pseudomonas aeruginosa. Environ Health Perspect 123: 749–758.

Fields BS, Benson RF & Besser RE (2002) Legionella and Legionnaires’ disease: 25 years of investigation. Clin Microbiol Rev 15: 506–526.

Galazzo G, van Best N, Benedikter BJ, et al. (2020) How to Count Our Microbes? The Effect of Different Quantitative Microbiome Profiling Approaches. Front Cell Infect Microbiol 10: 403.

Hamilton KA & Haas CN (2016) Critical review of mathematical approaches for quantitative microbial risk assessment (QMRA) of Legionella in engineered water systems: research gaps and a new framework. Environmental Science: Water Research & Technology 2: 599–613.

Hart ML, Meyer A, Johnson PJ & Ericsson AC (2015) Comparative Evaluation of DNA Extraction Methods from Feces of Multiple Host Species for Downstream Next-Generation Sequencing. PLoS One 10: e0143334.

Henne K, Kahlisch L, Brettar I & Hofle MG (2012) Analysis of structure and composition of bacterial core communities in mature drinking water biofilms and bulk water of a citywide network in Germany. Appl Environ Microbiol 78: 3530–3538.

Hermans SM, Buckley HL & Lear G (2018) Optimal extraction methods for the simultaneous analysis of DNA from diverse organisms and sample types. Mol Ecol Resour 18: 557–569. https://www.protocols.io/search?q=%22dna%20extraction accessed on 01-12-2023 p.^pp.

Hwang C, Ling F, Andersen GL, LeChevallier MW & Liu WT (2012) Evaluation of methods for the extraction of DNA from drinking water distribution system biofilms. Microbes Environ 27: 9–18.

Inkinen J, Kaunisto T, Pursiainen A, Miettinen IT, Kusnetsov J, Riihinen K & Keinänen-Toivola MM (2014) Drinking water quality and formation of biofilms in an office building during its first year of operation, a full scale study. Water Research 49: 83–91.

International Organization for Standardization (2017) Water quality - Enumeration of Legionella. Vol. ISO 11731:2017 p.^pp.

Kennedy NA, Walker AW, Berry SH, et al. (2014) The impact of different DNA extraction kits and laboratories upon the assessment of human gut microbiota composition by 16S rRNA gene sequencing. PLoS One 9: e88982.

Lazarevic V, Gaia N, Girard M, Francois P & Schrenzel J (2013) Comparison of DNA extraction methods in analysis of salivary bacterial communities. PLoS One 8: e67699.

Liu G, Zhang Y, van der Mark E, Magic-Knezev A, Pinto A, van den Bogert B, Liu W, van der Meer W & Medema G (2018) Assessing the origin of bacteria in tap water and distribution system in an unchlorinated drinking water system by SourceTracker using microbial community fingerprints. Water Res 138: 86–96.

Martin-Laurent F, Philippot L, Hallet S, Chaussod R, Germon JC, Soulas G & Catroux G (2001) DNA extraction from soils: old bias for new microbial diversity analysis methods. Appl Environ Microbiol 67: 2354–2359.

McMurdie PJ & Holmes S (2013) phyloseq: an R package for reproducible interactive analysis and graphics of microbiome census data. PLoS One 8: e61217.

McMurdie PJ & Holmes S (2014) Waste not, want not: why rarefying microbiome data is inadmissible. PLoS Comput Biol 10: e1003531.

Neu L, Proctor CR, Walser JC & Hammes F (2019) Small-Scale Heterogeneity in Drinking Water Biofilms. Front Microbiol 10: 2446.

Paranjape K, Bedard E, Whyte LG, Ronholm J, Prevost M & Faucher SP (2020) Presence of Legionella spp. in cooling towers: the role of microbial diversity, Pseudomonas, and continuous chlorine application. Water Res 169: 115252.

Paranjape K, Bedard E, Shetty D, Hu M, Choon FCP, Prevost M & Faucher SP (2020) Unravelling the importance of the eukaryotic and bacterial communities and their relationship with Legionella spp. ecology in cooling towers: a complex network. Microbiome 8: 157.

Pearman JK, Keeley NB, Wood SA, Laroche O, Zaiko A, Thomson-Laing G, Biessy L, Atalah J & Pochon X (2020) Comparing sediment DNA extraction methods for assessing organic enrichment associated with marine aquaculture. PeerJ 8: e10231.

Prest EI, Hammes F, Kotzsch S, van Loosdrecht MC & Vrouwenvelder JS (2013) Monitoring microbiological changes in drinking water systems using a fast and reproducible flow cytometric method. Water Res 47: 7131–7142.

Proctor CR, Reimann M, Vriens B & Hammes F (2018) Biofilms in shower hoses. Water Res 131: 274–286.

Props R, Kerckhof FM, Rubbens P, De Vrieze J, Hernandez Sanabria E, Waegeman W, Monsieurs P, Hammes F & Boon N (2017) Absolute quantification of microbial taxon abundances. ISME J 11: 584–587.

Rhoads WJ, Sindelar M, Margot C, Graf N & Hammes F (2022) Variable Legionella Response to Building Occupancy Patterns and Precautionary Flushing. Microorganisms 10.

Scaturro M, Chierico FD, Motro Y, et al. (2022) Bacterial communities of premise plumbing systems in four European cities, and their association with culturable Legionella. bioRxiv.

Shi Z, Kong Q, Li X, Xu W, Mao C, Wang Y, Song W & Huang J (2022) The Effects of DNA Extraction Kits and Primers on Prokaryotic and Eukaryotic Microbial Community in Freshwater Sediments. Microorganisms 10.

Stoeckel DM, Stelzer EA & Dick LK (2009) Evaluation of two spike-and-recovery controls for assessment of extraction efficiency in microbial source tracking studies. Water Res 43: 4820–4827.

Toplitsch D, Platzer S, Zehner R, Maitz S, Mascher F & Kittinger C (2021) Comparison of Updated Methods for Legionella Detection in Environmental Water Samples. Int J Environ Res Public Health 18.

Vosloo S, Sevillano M & Pinto A (2019) Modified DNeasy PowerWater Kit® protocol for DNA extractions from drinking water samples. protocolsio.

Wang X, Howe S, Deng F & Zhao J (2021) Current Applications of Absolute Bacterial Quantification in Microbiome Studies and Decision-Making Regarding Different Biological Questions. Microorganisms 9.

Watts SC, Ritchie SC, Inouye M & Holt KE (2019) FastSpar: rapid and scalable correlation estimation for compositional data. Bioinformatics 35: 1064–1066.

