## Supplementary figures and tables for "The impact of DNA extraction on the quantification of *Legionella*, with implications for ecological studies"

**Table of Contents**

**[Supplementary Figure 1. Impacts of pre-treatment steps on DNA extraction](#_Toc158985158)**

**[Supplementary Figure 2. Quantification of](#_Toc158985159) *[L. pneumophila](#_Toc158985159)* [via ddPCR](#_Toc158985159)**

**[Supplementary Figure 4.](#_Toc158985160)** **[Comparison of the quantification of](#_Toc158985160) *[Legionella](#_Toc158985160)* [spp. through ddPCR and Absolute Abundance](#_Toc158985160)**

**[Supplementary Figure 5. Positive control DNA extraction](#_Toc158985161)**

**[Supplementary Table 1. Primers, probes and ddPCR reagents and conditions for a duplex assay to detect](#_Toc158985162) *[Legionella](#_Toc158985162)* [spp. and](#_Toc158985162) *[L. pneumophila](#_Toc158985162)***

**[Supplementary Table 2. Primers, PCR reagents and conditions for the amplicon sequencing library preparation](#_Toc158985163)**

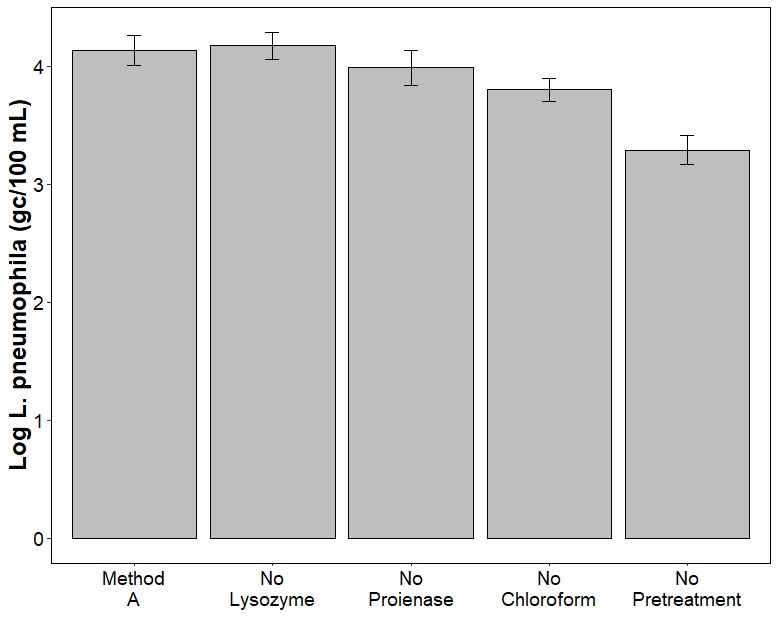

### **Supplementary Figure 1. Impacts of pre-treatment steps on DNA extraction**

The DNA extraction conducted in our laboratory, for both methods analysed, consisted of a few additional adaptations reported in section 2. The plot shows the impact of each of the pre-treatment steps on the quantification of *L. pneumophila* via ddPCR for the samples extracted with method A. The first bar (method A) represent the extraction carried with all the pre-treatment steps. The successive three bars indicate the impact of removing one pre-treatment step, while the last bar shows the impact of not performing any pre-treatment steps.

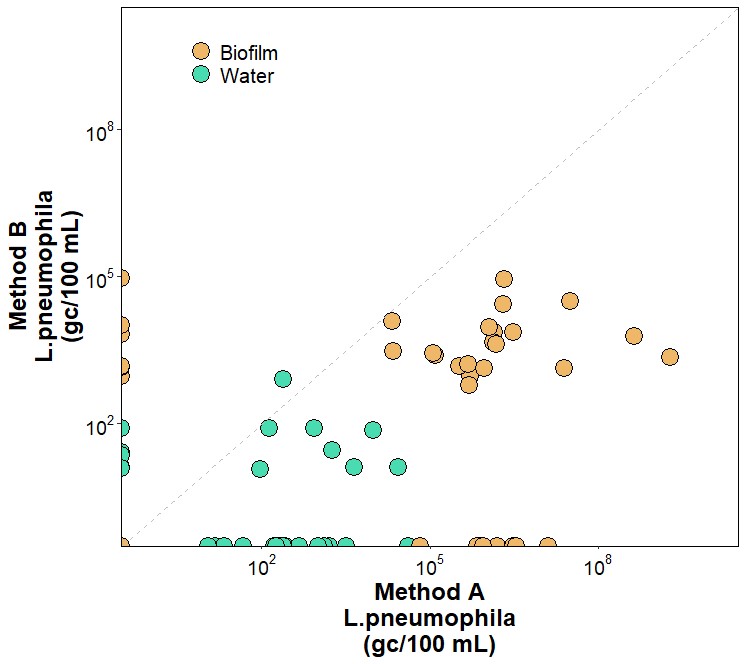

### **Supplementary Figure 2. Quantification of *L. pneumophila* via ddPCR**

The plot represents the quantification of L. pneumophila via ddPCR in gene copies per 100 millilitres. The quantification of the samples extracted with method B is indicated in the y-axis, while the quantification of the ones extracted with method A is indicated in the x-axis. Biofilm samples are indicated in orange, while water samples are shown in green.

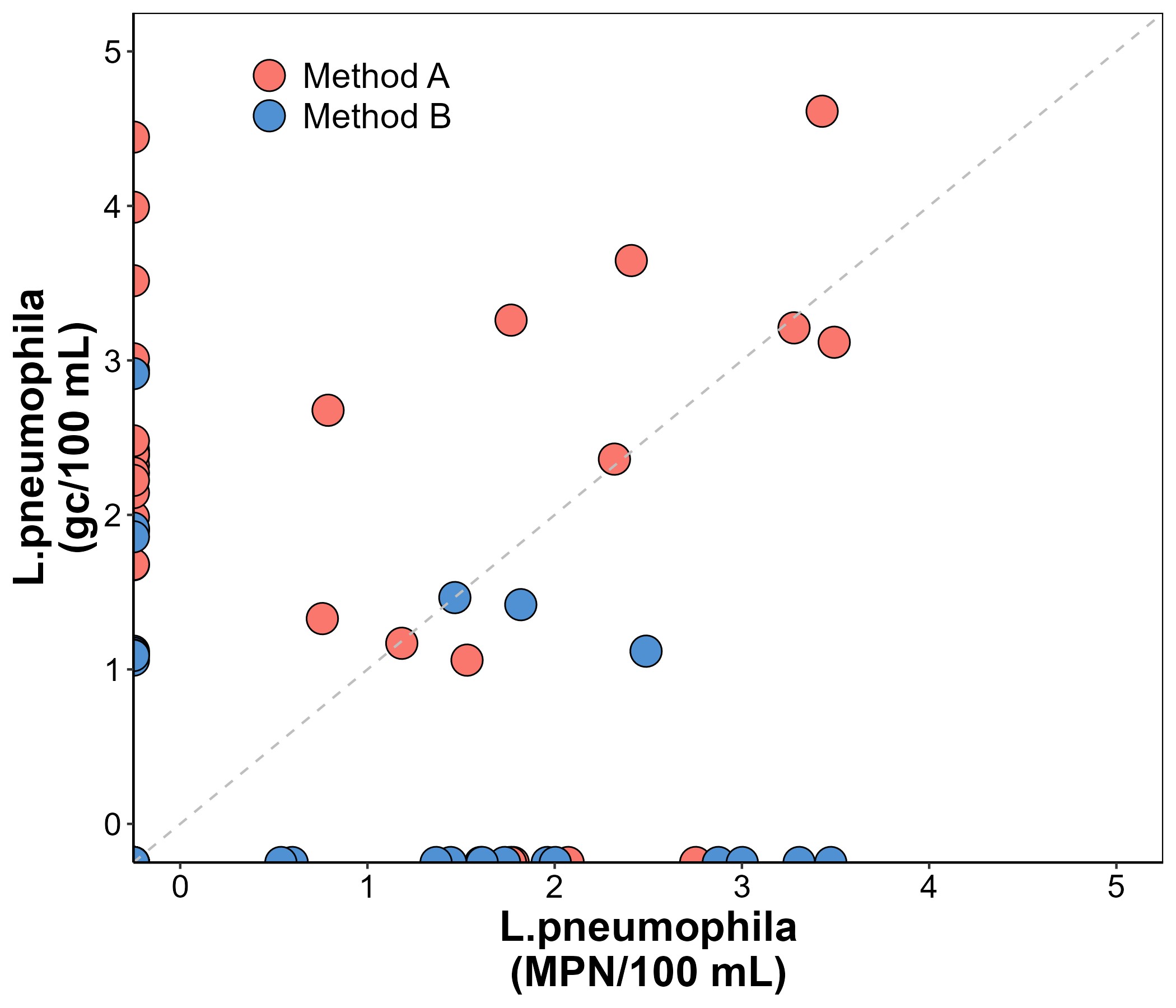

**Supplementary Figure 3. Comparison between ddPCR and cultivation-based quantification of *L. pneumophila***

The plot displays the comparison between the quantification of *L. pneumophila* through ddPCR (y-axis) and through culture-dependent method (x-axis). The samples are moreover separated by method: method A is represented in red, while method B in blue. The value are expressed as logarithm of the actual concentration measured in gene copies per 100 mL or Most Probable Number per 100 millilitres. *L. pneumophila* failed to be detected with ddPCR in samples extracted with method B in 10 samples that had positive cultures, highlighting the poor performance of the method. Method A and method B had positive *L. pneumophila* gene copy detections when culturable *L. pneumophila* was absent.

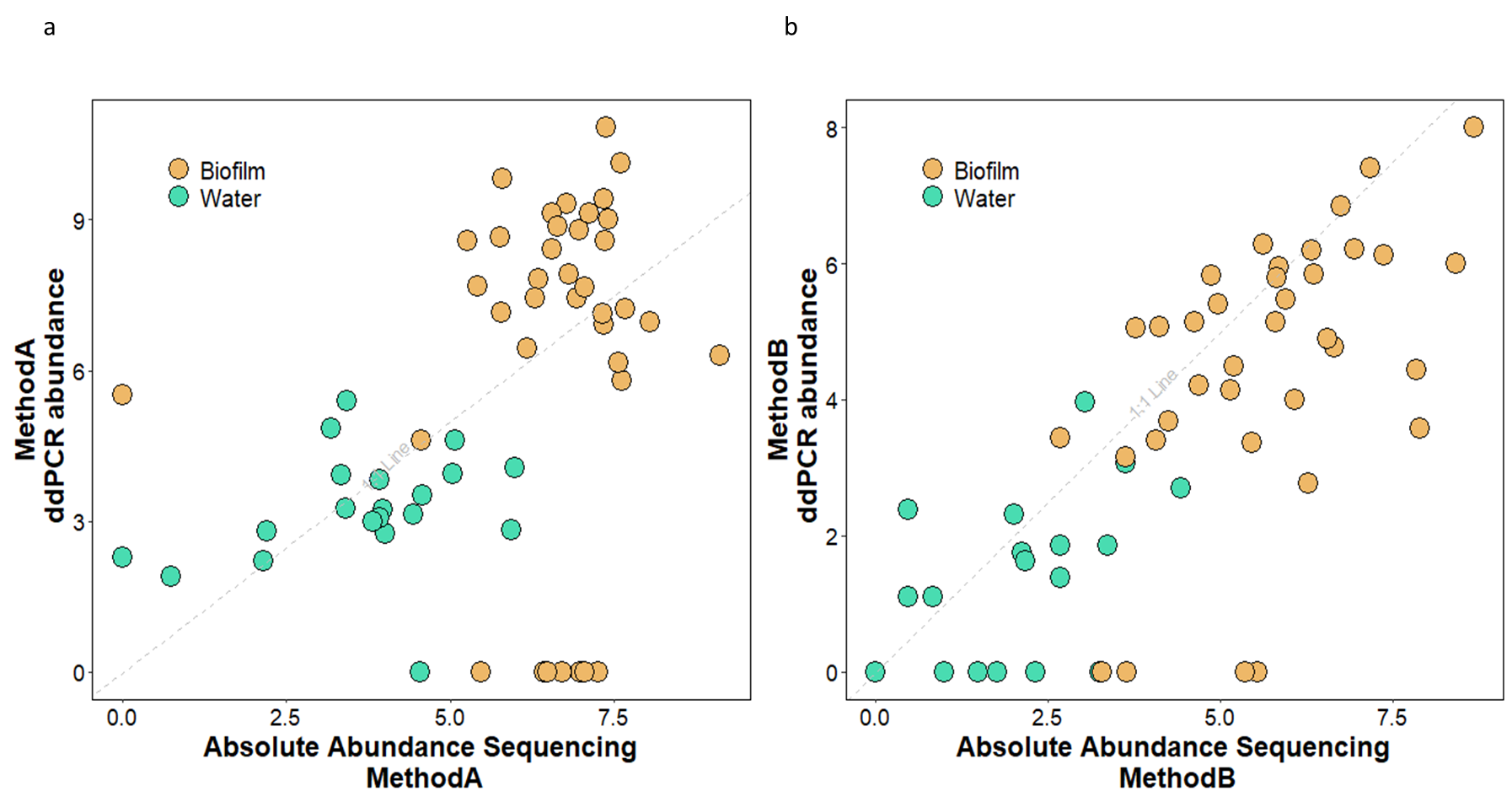

**Supplementary Figure 4.** **Comparison of the quantification of *Legionella* spp. through ddPCR and Absolute Abundance**

The plots show the comparison between the quantification of *Legionella* spp. via ddPCR (y-axis) and the Absolute Abundance calculated multiplying the 16S-amplicon-sequencing-derived relative abundance and the total cell count obtained via flow cytometry (x-axis). Plot A shows the comparison between the two quantification methods with respect to method A, while the comparison for method B is displayed in plot B. Biofilm samples are indicated in orange, while water samples are shown in green.

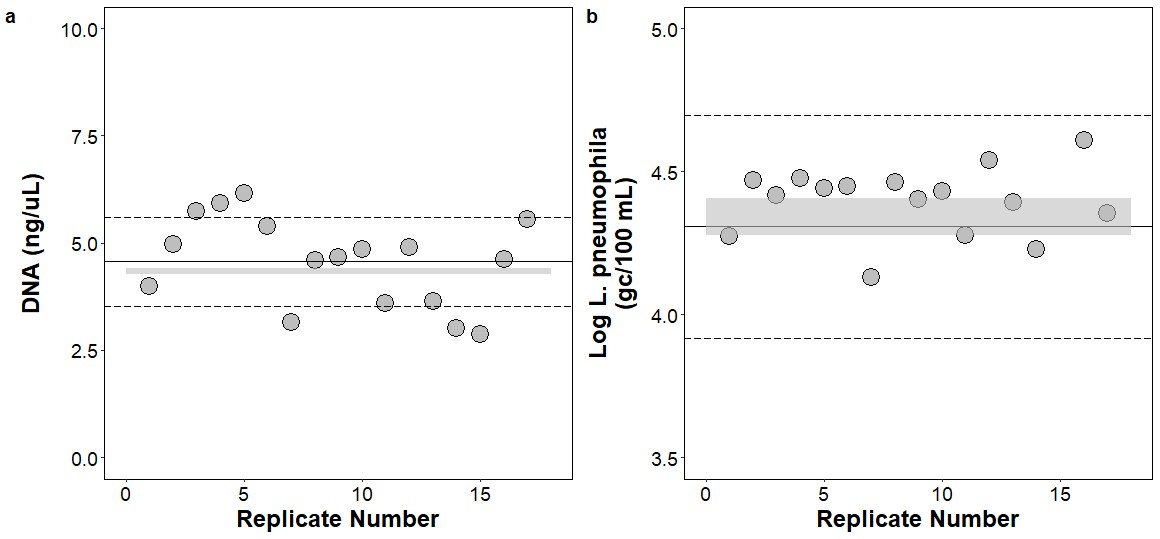

### **Supplementary Figure 5. Positive control DNA extraction**

The plots show the reproducibility of the DNA extraction process using several positive control prepared from the same environmental sample collected from a bioreactor in our laboratory. After collection, the sample has been filtered onto several filters in order to generate external positive controls to be extracted using the same procedure as for the samples, but taking into account the sample matrix effect.

Figure A shows the reproducibility of the extraction with respect to the quantification of the extracted DNA in nanograms per microliters, while figure B is relative to the quantification of *L. pneumophila* via ddPCR in gene copies per 100 millilitres. The continuous line represents the average value, while the dashed lines indicate the average value ± the standard deviation. The grey rectangle represent the expected values (expected DNA in figure A, MPN of *L. pneumophila* in figure B) ± the standard deviations.

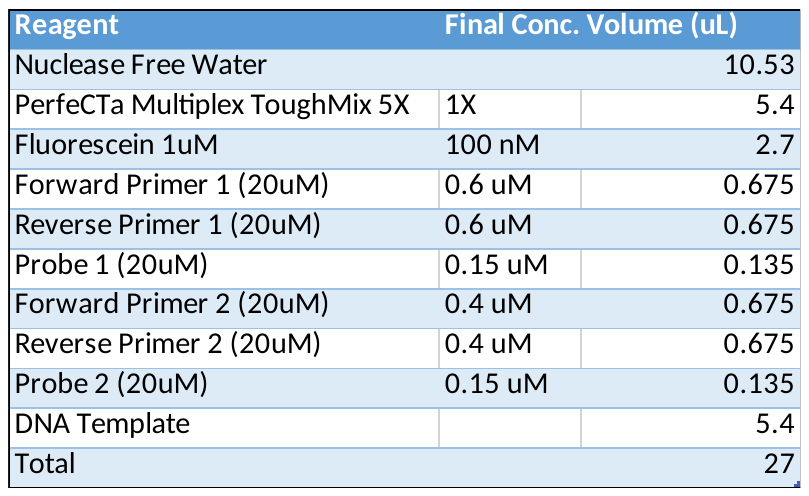

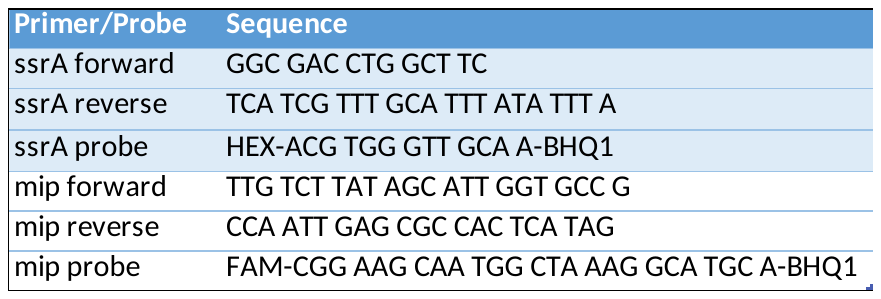

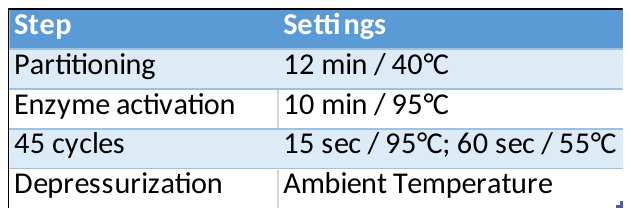

### **Supplementary Table 1. Primers, probes and ddPCR reagents and conditions for a duplex assay to detect *Legionella* spp. and *L. pneumophila***

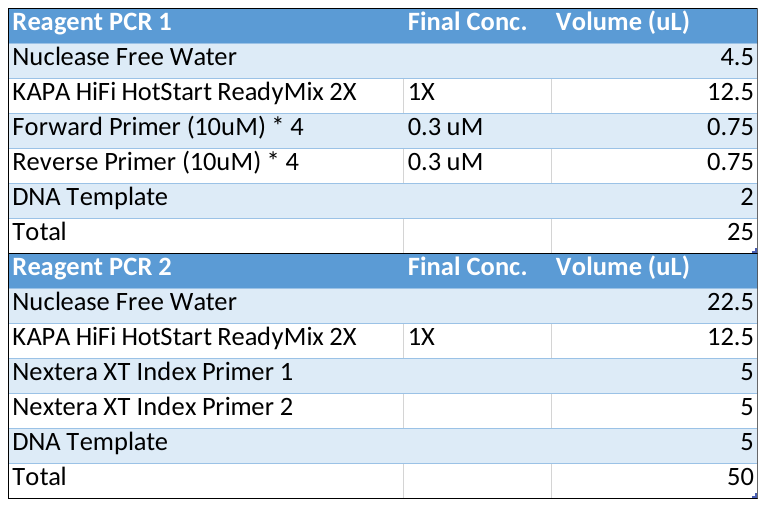

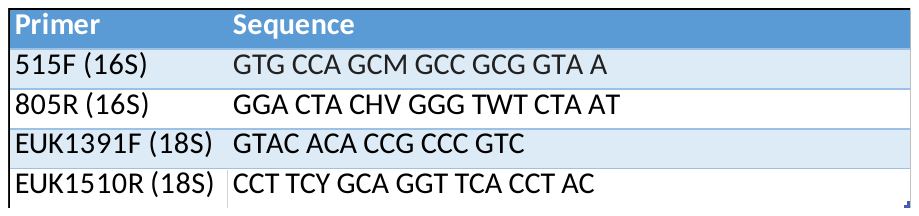

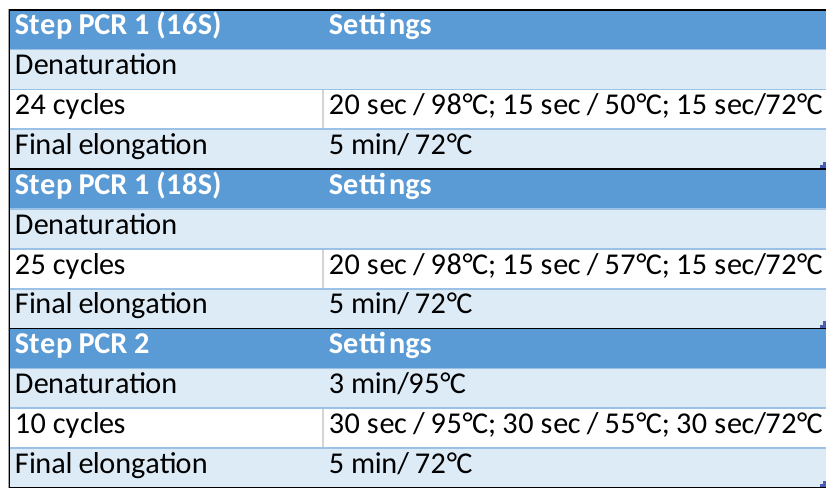

### **Supplementary Table 2. Primers, PCR reagents and conditions for the amplicon sequencing library preparation**
